# How animals endure by bending environmental space: Redefining the fundamental niche

**DOI:** 10.1101/2021.02.01.429132

**Authors:** Jason Matthiopoulos

## Abstract

For 70 years, the fundamental niche, the complete set of environments that allow an individual, population, or species to persist, has shaped ecological thinking. Yet, its properties have eluded quantification, particularly for mobile, cognitively complex organisms. Here, I derive a concise mathematical equation for the fundamental niche and fit it to population growth and distribution data. I find that, within traditionally defined environmental spaces, habitat heterogeneity and behavioural plasticity make the fundamental niche more complex and malleable, but also more predictable, than previously envisaged. This important re-evaluation restores the fundamental niche as a cornerstone of ecological theory and promotes it as a central tool for applied ecology. It quantifies how organisms buffer themselves from change by bending the boundaries of viable environmental space, and offers a framework for designing optimal habitat interventions to protect biodiversity or obstruct invasive species.

**ONE-SENTENCE SUMMARY:** The fundemental niche of animal species

## 1 Main text

Niche theory is as old as ecology^1^ and the concept of the fundamental niche^2^ holds prominence, well beyond the environmental sciences, in areas as diverse as evolutionary theory^3^ and cell biology^4^. The fundamental niche describes all environmental circumstances that allow a species to grow in numbers^2,5,6^ and is assumed fixed, for as long as the genotypic composition of the species stays the same^7^.

Like a Platonic ideal, the fundamental niche is never observed directly. Instead, its various manifestations, the realized niches, are observed in the distributions of species across landscapes^7,8^. Realized niches differ from the fundamental niche because the correspondence between habitat suitability and habitat occupancy is never perfect. Species are often absent from suitable habitat (due to dispersal limitations or exclusion by superior competitors) and present in unsuitable habitat (due to spillover of individuals from source habitats into neighboring sink habitats)^8^. Further, the fundamental niche may include habitats that are not currently present in geographical space.

The apparent separation of the fundamental niche from observed species distributions has led to recurrent debate about its utility and, even, calls for its abandonement^1,5,9–13^. However, although the fundamental niche is not essential for building descriptive models of where species are observed, it is indispensible for predicting where they could occur, outside our spatio-temporal frame of observation. Also, even when not explicitly acknowledged, the findings of studies on species’ global ranges, invasion potential, critical habitat and fine-scale habitat suitability, are interpreted in terms of *viability*, the defining notion of the fundamental niche^14^. Therefore, a quantitative understanding of the fundamental niche, is essential for transferrable models that can yield robust forecasts of species distributions, range expansions and extinctions^8,15–18^. “*The niche is here to stay*”^16^ because of our undisputed need to determine the viability of species in rapidly changing landscapes^10,19^.

Progress towards such understanding is obstructed by three knowledge gaps. First, despite a formidable body of conceptual and set-theoretic work that has detailed the distinctions between habitat suitability and observed species distributions^5,6,20,21^, we still require a general mathematical framework linking the fundamental to the realized niches. The indiscriminate use of the term “niche modeling” in *lieu* of species distribution modeling (SDM) has been counter-productive^14,22^ because it suggests that SDM, a method designed to map populations at pseudo-equilibrium (i.e. the realized niche), can quantify when small populations can grow exponentially (i.e. the fundamental niche). As long as the realized and fundamental niches are confused in this way, concepts that depend on them will also remain confused^23–25^.

Second, we have few^26^ statistical frameworks for estimating the fundamental niche empirically^1,8,19,27^, and none that specifically refer to higher animals. The assortment of heuristic methods, based on presence-only data, that have claimed to be able to estimate the fundamental niche without the need for data on population growth, in fact, estimate niche-related objects that are found “*at some unspecified point along a continuum between the fundamental and the realized niche*”^6^. Therefore, models that do not consider population growth or demographic rates, in addition to spatial data, are of limited use in an explanatory or predictive capacity.

Third, as a result of the above two gaps, we have only limited intuition about the shape and boundaries of the fundamental niche. Since its inception, it has been imagined as a closed and convex hypervolume in *n*-dimensional environmental space^2,23,27–30^. This image stems from the original descriptions of the fundamental niche in terms of morphological or physiological tolerances (e.g. temperature envelopes^15^), that define simple ranges along each niche dimension. However, it is becoming apparent that the structure of fundamental niches may be more complex^18,31^. In particular, it has recently been postulated that environmental heterogeneity and phenotypic plasticity make the boundary of the fundamental niche intractable^13^. This may be particularly the case for organisms that can sense heterogeneity and move selectively between different habitats.

Here, I address all three of these challenges. I derive a general, concise expression for the boundary of the niche that is mathematically tractable and statistically estimable from data. I apply this approach to compound spatial and temporal data from a passerine species and, using my results, I confront our current intuition about the fundamental niche. By doing so, I estimate the extent of endurance of plastic organisms, living in heterogeneous environments, in apparently inviable regions of niche space. I also discuss how this knowledge can help us solve seemingly impossible human-wildlife conflicts.

### 1.1 From *G*-spaces to *E*-spaces and back again

Geographical space (*G*-space) comprizes the three physical dimensions *x, y, z*. A location **s** = (*x, y, z*) in *G*-space may have *n* characteristics, such as scene-setting conditions (e.g. aspects of geomorphology, climate and soil composition), resources (e.g. amount of food, number of breeding sites) or risks (e.g. exposure to predators or pollution). These *n* variables form the dimensions of environmental space (*E*-space). Since Hutchinson^2^, this is also known as “*niche space*”, because he envisaged the fundamental and realized niches as portions of *E*-space. A point **x** = (*x*_1_, …, *x_n_*) in *E*-space uniquely defines a local environment, or habitat^32^. Within any given landscape, each habitat has an associated availability value (e.g. the total area of that type of habitat across *G*-space). I will denote availability by *f*_**x**_, below. Fig. 1 visualizes the correpondence between *G* and *E* spaces and prompts three important points. First, several landscapes can be constructed by re-arranging the same ingredients in *G*-space (e.g. Figs 1a and b). Second, the shape of habitat availability clouds in *E*-space can be very complex^24^, even including discontinuities and holes^31^. Third, availability plots in Hutchinson’s *E*-space hold no information about the spatial contiguity of habitats, and are therefore unable to communicate the *context* in which an organism finds itself. Hence, although only one *E*-cloud can be constructed from a given *G*-landscape, infinite landscapes can be constructed from an availability cloud in *E*-space^33^ (the transition from *G* to *E* is irreversible).

**Figure 1:**
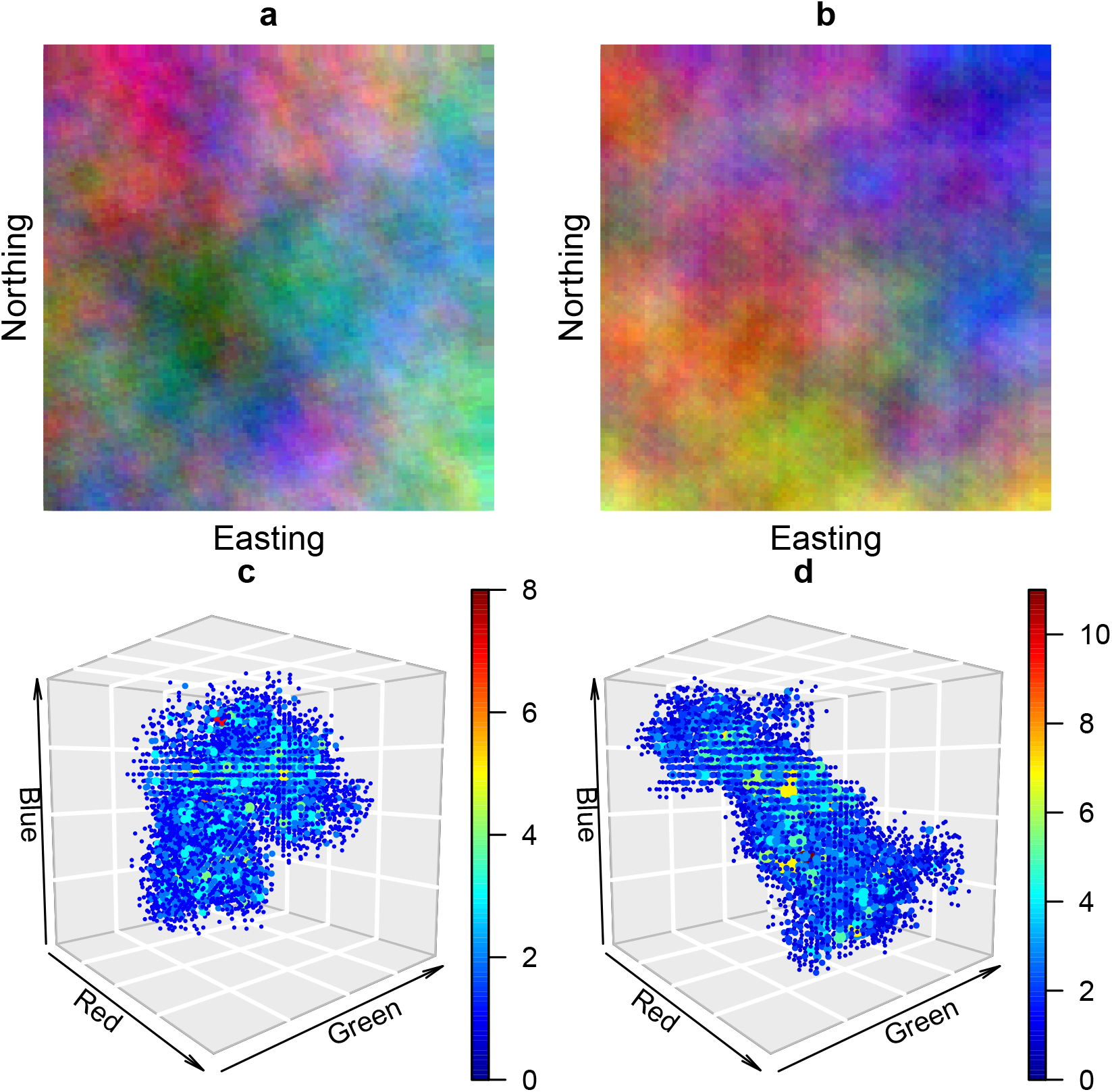
Illustration of Geographical (top row) and Environmental (bottom row) spaces. A *G*-space is a landscape, here mapped in the two dimensions of Easting and Northing. Two such landscapes (plates a & b) were created by mixing similar total amounts of three environmental variables. Each enviromental variable is represented by a primary color red, green or blue. As different intensities of the three primary colors overlap locally in plates a & b, they create a wide variety of mixture colors, each symbolizing a different habitat. The three axes of *E*-space (plates c & d) represent these three environmental variables (primary colours). Each mixed color in *G*-space (a unique habitat type) is a point in the 3-dimensional *E*-space, characterized by its constituent primary colours. Recording a dot at a particular point in *E*-space implies that the corresponding habitat can be found within the landscape in *G*-space. The new colour heat-scale in *E*-spaces c & d has a distinctly different interpretation. It represents the availability of each habitat type, the frequency with which this exact combination of environmental values occurs in each of the two landscapes above. The more spherical shape of the availability cloud in plate c, as compared to plate d, is entirely the result of chance. More symmetric clouds than c, and more irregular clouds than d are possible.

Traditionally^2,23,27–30^, niches have been imagined as subsets of *E*-space, objects not dissimilar to the 3D cloud in Fig. 1c. Indeed, if an animal inhabiting Fig. 1a divided its time equally between all points of that landscape, its realized niche would coincide with Fig. 1c. However, there has always been confusion between the fundamental niche as a subset of *E*-space and as a place, in *G*-space^23^. This lingering confusion stems from the tenable notion, that a point in *E*-space cannot be considered in isolation from its geographical context^13^. For instance, a situation not currently captured by the niche literature is that animals are routinely able to survive in niche spaces where no *single* point is sufficient for their survival and reproduction (I will tentatively call this the “zero niche paradox”, a potential focus for future methodological work, see Methods). Often, vital resources are mutually exclusive in space (e.g. at a fine spatial scale, water holes and grazing land cannot coincide), and animals have to perform short commutes or longer-range migrations to satisfy all their life-history requirements. By moving across heterogeneous landscapes, animals can experience different types of habitats, and by actively selecting to use some over others, they demonstrate high levels of behavioural plasticity. If we imagine that the two landscapes in plates a & b of Fig. 1 happen to be the home ranges of two animals from the same species, then habitat suitability and the resulting viability of these two animals may differ, even though they have access to the same average amounts of resouces. So, we must consider the animals’ fitness in the light of their *entire* environmental profile (i.e. the whole clouds in Figs 1c, d), not merely any single point in *E*-space.

It may be argued that this is a problem of scale^34^ and that summarizing (e.g. averaging) the environmental variables at spatial resolutions comparable to the mobility of individuals may restore the fundamental niche to its classic form (i.e. mapping viability to single points in *E*-space). However, as can be seen in Fig. 1, even though the average amounts of resources available to two organisms may be similar (i.e. the total amounts of red, green and blue used to paint the landscapes in Figs 1a,b), their particular configurations into habitats may be very different. Spatial and temporal heterogeity will generally affect the viability of an organism^20^ so it is not clear whether the habitat homogeneity implied by averaging environmental variables at coarse spatial scales still permits us to correctly represent species responses to habitat.

It is important therefore to ask whether the existence of such heterogeneities, and the complex responses of animals to them, require us to modify our intuitive understanding of the fundamental niche. Does the ubiquitous fact that animals exploit environmental heterogeneity matter for the size, shape and predictability of their fundamental niche? Below, I find that the answer is yes, even in very limited environmental spaces, using the most rudimentary levels of environmental heterogeneity. I first present the niche boundary equation and then its application to a particular species, the house sparrow, leaving the full derivation and analysis details for the Methods.

### 1.2 General definition of the fundamental niche

Hutchinson envisaged the fundamental niche as the set of points **x** in *E*-space yielding non-negative intrinsic population growth^5,6^, allowing a population of genetically identical organisms to invade and occupy these habitats^6^. Habitat can be connected to the growth in population size *N_t_* by the average, per-capita fitness *F* (**x**), associated with each point in *E*-space^35^

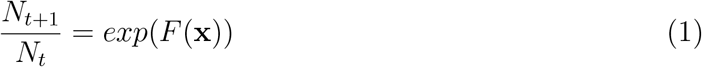

under the assumption that the population is sufficiently small for crowding effects to be negligible. Then^19^, the fundamental niche is the *E*-space subset,

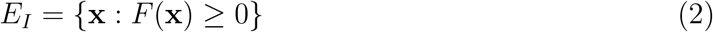

and the niche boundary is obtained by setting *F* (**x**) = 0.

However, viability does not only depend on habitat **x**. Spatial heterogeneity may offer animals different options within their accessible space. Then, **x** becomes the central vantage point from which they see this heterogeneous landscape. For example, **x** may be the habitat characterizing the current location of a nomad in the landscape, or the habitat at the centroid of the home range of a central-place forager. Alternatively, it may be a summary of an animal’s surroundings, such as the average habitat within its home range. Ultimately however, **x** is a reference habitat in *E*-space, around which we want to evaluate viability, in order to determine whether **x** belongs to the fundamental niche, or not. Given a reference habitat **x**, the availability (*f*_**z**|**x**_) of other habitats **z** in the surrounding landscape will depend on spatial structuring and the mobility of the organism^33^. From this palette of available habitats, animals choose to use some more than others. Habitat preference (*w*_**z**|**x**_) can capture such variations in usage^32^. Finally, each minute of usage of each available square meter of a habitat will contribute an incremental amount *F*_**z**_ to that animal’s overall fitness^35^. In the Methods, I show that the boundary of the niche can, in general, be described by the equation

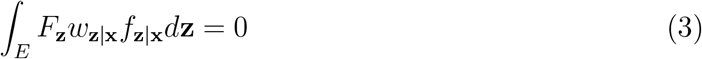

The integral is proportional to the fitness *F*(**x**) around a reference habitat **x**. It aggregates all nearby habitat-specific contributions *F*_**z**_, weighted by the availability (*f*_**z**|**x**_) of each habitat and by preferrential usage (*w*_**z**|**x**_) of habitats by members of the species. It has been shown^32,33,35,36^ that all the components in this expression can be estimated from data. In particular, the formulations of *f*_**z**|**x**_ can describe much more complex availability clouds in *E*-space, than those shown in Figs 1c,d, but can also represent the contiguity and structure of habitats in *G*-space^33^. In addition, *f*_**z**|**x**_ contains information about the mobility of the organism, enabling it to capture the scale at which a typical individual experiences the ambient heterogeneity in its environment. Implicit information on these two quintessentially geographical properties (habitat contiguity and animal mobility) allows the integral over *E*-space in eq. (3) to capture the habitat context around **x**. Although originally proposed to account for spatial structuring, this model can readily be extended to temporal structuring (e.g. trend, stochastic or seasonal components), hence extending the notion of viability to temporal change^18^, accounting for aspects of plasticity displayed by sessile organisms, such as plants.

The solutions of eq. (3) are not merely points in *E*-space, but fully parametric descriptions of entire landscapes, that can offer the species neutral fitness. The equation has infinite solutions in its extended parameter space, which is much higher-dimensional than *E*-space. These extra dimensions arise from the need to describe complex habitat availability distributions in *E*-space (using parameters to capture higher moments of resource distributions, such as variance, skewness, outliers, but also multimodality). For example, for an *E*-space of *n* orthogonal environmental variables, allowing a trimodal marginal distribution of availability in each environmental dimension, results in a fundamental niche space of at least 4*n* dimensions (characterizing the positions of the three modes and an identical variance around each mode). Even in the case of unimodal availability (corresponding to *n*-d elipses in *E*-space), describing a heterogeneous environment requires twice as many dimensions as Hutchinson’s niche space. Only completely homogeneous environments can be sufficiently described by *n* environmental dimensions. Although impossible to visualize, solutions to this equation are entirely possible to retrieve. As will be seen in the house sparrow example below (see eqs (25) and (26)), for some natural histories, it may even be possible to describe the fundamental niche using simple integral-free algebraic expressions. If the objective is not to exhaustively describe the set of solutions, but rather, to examine if a particular landscape is inside or outside the niche^21^, then numerically calculating the (fully parameterized) integral in eq. (3) will give positive or negative values respectively.

### 1.3 The fundamental niche of house sparrows

House sparrows are a good species for niche studies^36,37^. They have a wide distribution, tempered by global climate patterns^37^, but their viability is affected by microhabitat composition^36^. They are considered invasive pests in the Americas, but threatened endemics in Europe. They are also greatly influenced by lansdscaping trends and urbanisation, making them exemplars of human-wildlife conflict.

The particular study in^36^ looked at fine-scale suburban garden composition within the home ranges of different sparrow colonies. Home range composition was described in terms of six land cover variables. Sparrows were not particularly selective within their home ranges (the habitat preference explained only 33% of observed patterns of usage), but the population models based on habitat availability and usage captured 81% of the variability in colony growth rates, under cross-validation. This high predictive ability was achievable with detailed information on the distribution of three of the six variables (percentage of grass, bush and roof structures). Superabundance of these variables was detrimental to population growth. Sparrow colonies were least tolerant of high percentages of lawn and performed better in the presence of bushes and roof structures.

In eqs (20), (23) and (24) of the Methods, an integral-free expression is derived for house sparrows, describing their fundamental niche’s boundary in terms of garden composition and heterogeneity therein, fitness parameters and habitat selection parameters. Although only three environmental variables were included in the model, a total of 57 dimensions were used to describe the heterogeneity in garden compositions.

To explore how a high-dimensional representation of the niche would behave once projected onto 3-d Hutchinsonian *E*-space (Fig.2), I explored the viability of all home ranges with feasible habitat compositions. Feasibility required that the sum of land cover variables (at any point of the home range) could not exceed 100%, a constraint represented by the semi-transparent inclined plane in the four panels of Fig.2. I specified the fully-fitted model to the simplest scenario of habitat heterogeneity, using unimodal Gaussian spheres in *E*-space. This meant that the niche model was explored in 4-d space (i.e. the three mean values for the land cover variables with the addition of a common variance dimension describing habitat heterogeneity within a home range). The extent and consequences of this form of heterogeneity in the sparrows’ home ranges could then be examined by increasing a single variability parameter *σ* from zero (i.e. homogeneous home range), to its largest feasible value. First, under the assumption of home range homogeneity (i.e. under the Hutchinsonian *n*-hypervolume definition of the fundamental niche), I found that the niche occupied 15% of feasible *E*-space (Fig.2a). Introducing habitat heterogeneity and sparrow selectivity allowed an inflation of the complete set of habitats that could be included in viable home ranges to 25% of feasible space (Fig.2b). Improvements in the maximum fitness of animals encountered in different habitats, occurred primarily in the centre of viable space (Fig.2c) because this form of spherical heterogeneity could not be high close to the borders of the feasible niche space. The fitness characterizing entire home ranges was also improved compared to the homogeneous scenario (Fig.2d), because sparrows could select to use the better-than average parts of their heterogeneous home ranges. Heterogeneous home range fitness improved in 9% of the previously viable homogeneous home ranges and an additional 0.12% of previously inviable ranges bacame viable. The average improvement in fitness was 7.93% (95CI: 0.04%,50.74%). It is notable that these levels of niche and viability inflation are observed in a model with only three environmental covariates, under the most rudimentary form of heterogeneity (i.e. a shared variance parameter for all environmental variables, with no multimodality or asymmetries) and in a study species that demonstrated limited selectivity for habitats within the individual colony range. This is therefore a stringent test of the original hypothesis, that heterogeneity and behavioural plasticity alter the shape of Hutchinson’s fundamental niche in a real species.

**Figure 2:**
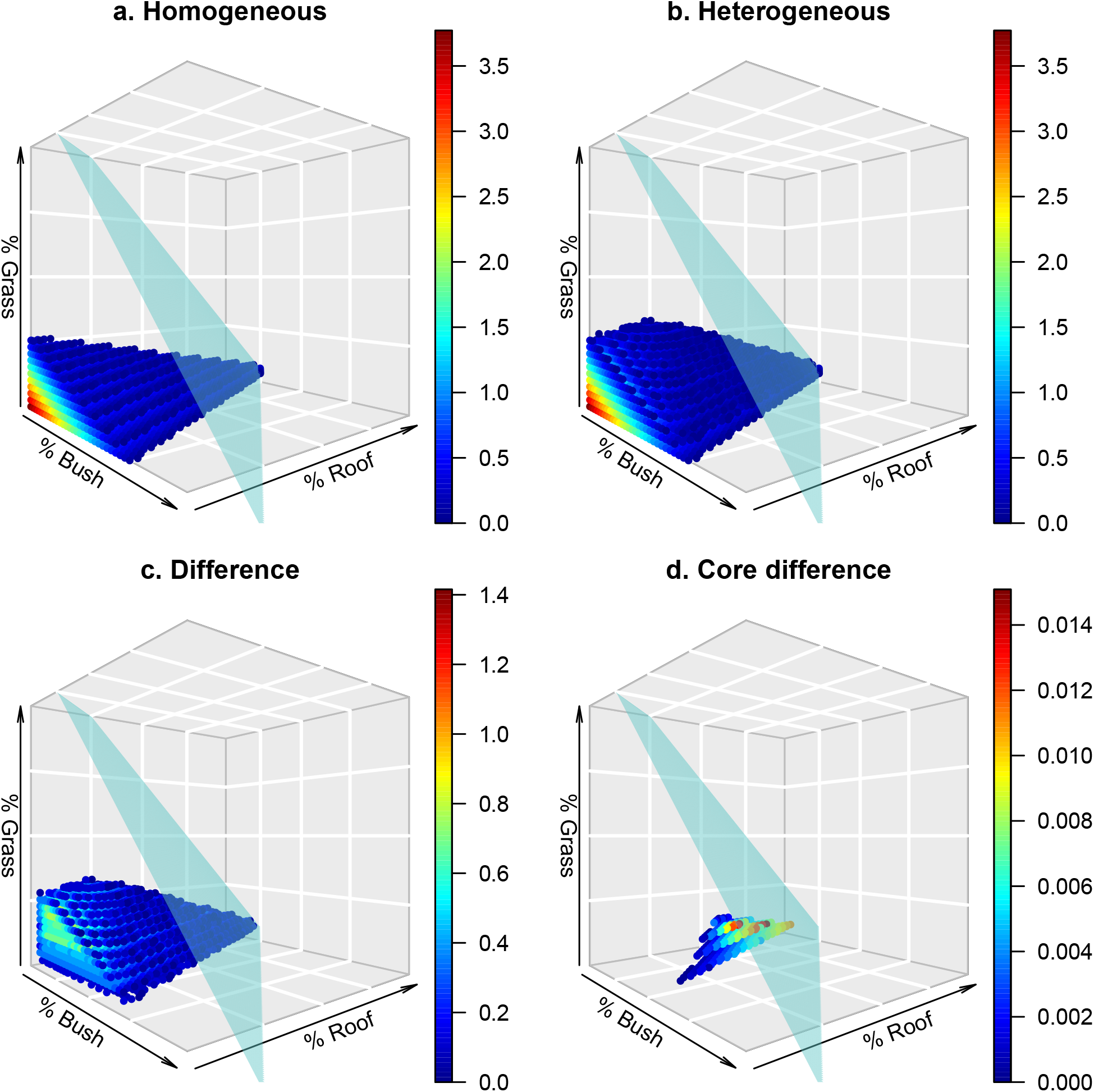
Visualizing the fundamental niche of house sparrows in homogeneous and heterogeneous home ranges. The transparent plane represents a feasibility boundary, preventing the sum of the three land cover variables from exceeding 100%. The three axes are plotted from 0% to 80%, to better focus on the viable region. Relative fitness values (or their differences) are colored on a gradient from high (red) to low (blue). The homogeneous scenario (a) assumes that all parts of a home range have the same composition as the average of the home range - the set of coordinates defining a point in feasible space. This is therefore a data-driven representation of Hutchinson’s *E*-space hypervolume for sparrows. The heterogeneous scenario (b) encloses all habitats that occur in viable home ranges when the heterogeneity in a home ranges maximizes fitness. The difference (c) between the predictions of the two scenarios highlights which regions of the homogeneous fundamental niche benefit from the existence of habitat heterogeneity and sparrow selectivity. The core difference (d) compares the change in relative fitness for homogeneous and heterogeneous home ranges with the same average habitat composition

### 1.4 Discussion

I have shown that the fundamental niche can be formulated mathematically and its parameters estimated empirically from distributional and population growth data. The key to achieving this is to treat the multiplicity of realized niches as an inferrential strength, rather than a nuisance. It is true that the fundamental niche of a species cannot be estimated from a single snapshot of its population distribution^1,13^. However, I show that multiple realizations, when treated as sampling instances in statistical estimation, allow the partly obscured picture of the fundamental niche to be assembled from different viewpoints. In this way, the niche object, estimated from case studies such as the sparrow example, is an approximation of the species’ fundamental niche that will asymptotically improve as more (contemporary and historical) data from the worldwide range of a species are included in the modelling. The rate at which this approximation converges with diverse data, for different taxa, is an important future research question.

My key theoretical finding, that for a given number of environmental dimensions, the niche is a more malleable volume than has thus far been imagined, has important practical implications. It suggests that niche boundaries in *n*-dimensional hyperspaces appear “fuzzy”^13^ not just because of stochasticity, but because they exist in much higher-dimensional spaces. This implies that the niche is more deterministic (hence, predictable) than we might conclude by viewing purely its low-dimensional projections. Heterogeneous environments can be more favorable to a mobile and selective species than homogeneous ones (even ones that are **on average** marginally better) and the framework presented here can quantify exactly how bufferred real animals are from hostile environments.

This re-evaluation of the dimensionality of the niche means that exhaustive visualizations are not possible, even for the simplest of examples. The image in Fig. 2b, is a 3-d projection of a 4-d model, which represents a gross simplification of a 57-d model of the sparrow niche, expressed solely in terms of land-cover). However, having a compact and numerically efficient expression that describes the boundary of the niche is arguably more useful than visualizing it, because it allows us to identify viable environments^21^ and to manage land cover so as to optimize population viability. By subtly modulating the availability and heterogeneity of environmental variables, this framework allows us to engineer viability for a species where there previously may have been none^36^. As Fig. 1 illustrates we can do this without necessarily changing the overall proportions of habitats in a landscape, by re-arranging them spatially. Given the reality of conflicts between conservation, resource/pest management, wealth creation, food security and complex ecosystem dynamics, where the overall amounts of land cover for each activity are often fixed, achieving such accurate mitigation will prove invaluable in the future.

With these priorities in mind, the road ahead for developing the tools of niche theory seems both clear and open. Progress primarily demands the convergence between mechanistic and statistical approaches. Mechanistic approaches propose to build fundamental niches from the ground up, using only biological first principles^15^. This emphasis on mechanism faces the challenges of reductionism^1,20,26,38^ and currently seems restricted to capturing well-understood and univariate physiological tolerances (e.g. thermal envelopes). Nevertheless, it is correct that statistical analyses of the niche cannot go far without biological mechanism^1^. Spatial data cannot estimate the niche without information on population growth, density dependence and demography. The models used in this paper were asked to predict the intrinsic growth rate of the population^16^, but they were fitted by accounting for density dependent effects, both in distributional^32,35^ and growth data^35,36^.

Increasing the mechanistic content of the models presented here could also deal with two closely related limitations of niche models^11^, their apparent inability to deal with resource depletion (and broader intraspecific effects) and the difficulty in capturing interspecific interactions such as predation, mutualism or competition. Both of these require us to think of the niche in more dynamic terms^5,16,26^ and, potentially, to arrive at a mathematical formulation that hybridizes the Grinellian & Eltonian ideas of the niche as a *role in the ecological community* with the Hutchinsonian idea of the niche as a *volume in E-space*^5,6,11^.

Such a formulation would capture dynamical interactions by including other species (prey, predators, competitors) as additional dimensions of *E*-space^8,23^, an idea that may not have universal appeal because it integrates the niche with its community context. However, as shown here, it is both possible, and necessary to formulate the fundamental niche in a context-dependent way, as long as all possible contexts are considered. Indeed, the idea of simultaneous modelling of the niches and distributions of multiple interacting species is already two decades old^39^ and rapidly gaining momentum^19,40,41^.

A final important extension of this modelling would be the inclusion of individual variation^13^. Much of the material presented here was developed for populations or species, but has continuously referred to the mobility and behaviour of “typical” individuals. Therefore, any predictions from these models do not capture potential phenotypic variation that could, in principle, characterize each sparrow by its own, individual fundamental niche. This hierarchy of fundamental niches, from individuals, to populations, to entire species has important implications for our understanding of evolutionary processes^3^ in dynamic landscapes. Versions of the model presented here, extended to include individual variation, are entirely feasible, given the computational efficiency of the algebraic calculations involved. The fundamental niche is the phenotypic link on which natural selection operates^18^ and therefore, a good understanding of evolutionary processes at the level of species-habitat-associations can only benefit from knowledge of the shape and properties of the fundamental niche^18^.

### 1.5 Conclusion

As ecology is moving towards transferrable models of population viability and distribution^10,17,42^, the fundamental niche is an indispensible concept^14,16^. The niche must be pattern- and data-driven^14,26^, but it must be rooted in ecological principles and applied imperatives^12^. I have argued for the use of hybrid^11^ correlational-mechanistic approaches^15,26^, that bring to the fore issues of population growth, transient dynamics, resource depletion and inter-specific interactions. Taking environmental heterogeneity and phenotypic plasticity^13^ into account has led to a re-evaluation of the complexity of the fundamental niche, a route towards its evolution as a concept, rather than its abandonement. These results carry optimistic messages about the resilience of animal species to anthropogenic change and the room for manoeuver in mitigating human-wildlife conflicts.

## 1.6 Acknowledgments

Earier drafts benefited from feedback by Daniel Haydon, Fergus Chadwick, John Harwood, Geert Aarts, Ewan Wakefield, Neil Metcalfe, Luke Powell, Pat Monaghan and Ross MacLeod. The IBAHCM provided the academic environment and time that made this work possible. ** Author contributions** JM conceived the mathematical framework, conducted the analysis and wrote the paper. ** Competing interests** None.

## 2 Methods

### 2.1 General definition of the fundamental niche

Theories of the fundamental niche envisage a mathematical connection between environmental space (*E*-space) and the viability of a species^5,6^. Each of the dimensions of *E*-space measures a biotic or abiotic environmental variable, i.e. a continuous, discrete or qualitative random variable representing a condition (e.g. pH, temperature, altitude), resource (e.g. soil nutrients, prey, breeding sites) or perceptible threat (e.g. predators, pollution). A point **x** in *E*-space can be conveniently referred to as a habitat^32,43,44^. The availability *f*_**x**_ of a habitat **x** is difficult to conceptualize and quantify, because it depends on how finely habitats are defined and how accessible they are from the vantage point of the study organism. To begin with, we can think of availability as the amount (e.g. total area) of a habitat that is accessible to an individual or a group. This definition makes two implicit assumptions. First, that habitats in *E*-space are finite (not infinitesimal) volumes, so that it makes sense to measure the area they occupy in *G*-space. Second, that any point in *G*-space is either fully or not at all accessible by the organism. Neither of these assumptions imply loss of generality in what follows, because relaxing them leads naturally to infinitesimal definitions of habitat where availability is a probability density corrected for accessibility. Therefore, the availability scalar field **f** is a probability density function that assigns a value *f*_**x**_∈ [0, 1] to each point **x** in *E*-space. For a given organism, or group, we assume that *∫_E_ f*_**x**_*d***x** = 1.

For the purposes of species- or population-level models, the concept of individual fitness (as a characteristic of a genotype or phenotype) is often generalized to populations, by linking the average fitness *F* of genetically or phenotypically similar individuals to population growth, using general models of the form (e.g. eq. (3.9) in^45^)

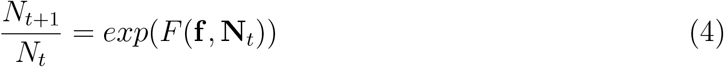

in which the population’s fitness is determined by environmental (**f**) and density dependent (**N**_*t*_ = {*N_t_, N_t−_*_1_, …}) influences. Individual variation in genotypes and phenotypes within a population implies that eq. (4) is merely a deterministic model for the mean of a distribution of individual fitnesses within a population. This ecological use of average fitness has been extensively discussed in the literature^46–50^ and eq. (4) has a long history of use in evolutionary models^51–53^. It also specifies a mathematical link between the environment of a species and its ability to grow, hence allowing us to formalize the concept of the niche at a species level, rather than the level of the individual. To adhere to Hutchinson’s original definition of the fundamental niche we must simplify this model to the intrinsic growth rate (i.e. a density independence scenario, where **N**_*t*_ ≈ **0**), and the experience of a single habitat (either because the whole of G-space is homogeneous, or because the organism is sessile).

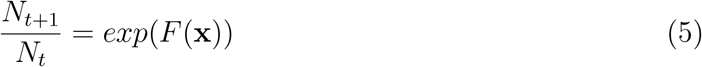

Then, the *fundamental niche* is a region in *E*-space that can be invaded and occupied by a population of genetically similar organisms^6^. More formally, if **x** is a point in *E*-space and *F* (**x**) is the fitness associated with that point, then the niche is the set

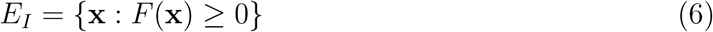

However, a number of specific challenges arise when we try to connect this definition, in *E*-space, with what happens in geographical, or *G*-space. Making this connection is important because data on habitat variables and observations of animal use are collected in *G*-space. As has been argued previously^33^, information is generally lost when transferring geographical data into *E*-space. These challenges are partly responsible for the difficulty in estimating fundamental niches from wildlife and environmental data.

### 2.2 Animal mobility and the niche

The main challenges in formalizing the niche concept for animals, is that they are predominantly mobile organisms and that, even the more primitive animals display complex capabilities of perception and cognitive control of their movements, such that they can actively select which habitats to use. This requires us to consider just how mobile an animal is (over a given time scale) and what habitats are likely to be available to it from its position. From the perspective of *E*-space, mobility determines the capacity of an organism to reach and use different habitats **z** given that it is currently occupying a particular habitat **x**, but this effect can only be captured by considering habitats in *G*-space. If, at a set of geographical coordinates **s**, an animal encounters the habitat **x**, we may be able to anticipate the types of habitat that are likely to be available to it, within its range of accessibility^33,54^. Therefore, an important aspect of the fitness *F* (**x**) attained by an organism when it is located at a particular habitat **x**, at a set of spatial coordinates **s**, is that it depends on context^13^, i.e. the habitat composition in the neighborhood of those coordinates. Because of this, it is necessary to distinguish between the context-specific version of fitness *F* (**x**) an d the unconditional fitness contribution of a single habitat, *F*_**x**_, defined here as the long-tern fitness characterizing a completely *sessile* individual constrained within habitat **x**. For example, with most plants and the sessile stages of some animals (e.g. porifera and anthozoa), we can assume that *F* (**x**) = *F*_**x**_. Similarly, when the mobility of an animal is small (e.g. echinoderms) compared to the heterogeneity of their environment, *F* (**x**) ≈ *F*_**x**_. Conditional usage (here, denoted by *u*_**z**|**x**_) can help express a relationship between conditional and unconditional fitness:

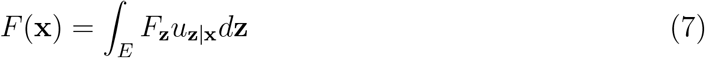

The integral gives overall fitness *F*(**x**) at **x** as the usage-weighted average of habitat-specific fitness contributions *F*_**z**_, of all habitats across *E*-space. We consider unconditional fitness and conditional usage in turn. Unconditional fitness can be adequately represented by quadratic polynomials of habitat variables^35,55^. Specifically, linear (hence, monotonic) terms may be used to describe responses to environmental resources and risks while parabolic terms can describe peaks in the responses to conditions. Two distinct components of fitness have been considered^35^, with and without the effect of density dependence. Although the density dependent component must be included when fitting to real population data, in defining the fundamental niche we are interested in the population’s intrinsic growth rate which can be obtained from the density-independent part of the model (assuming no Allee effects are in operation at small population sizes).

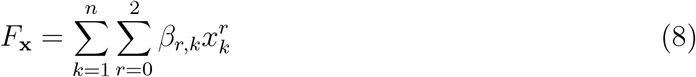

where the first sum is over *n* habitat variables and the second sum generates the quadratic polynomials with terms of order *r*.

The conditional habitat usage component *u*_**z**|**x**_ in eq. (7) may be expressed as a function of habitat preference and habitat availability. This approach is taken by a broad class of methods under the general name of Habitat Selection Functions (HSFs), a term previously introduced^35,56^ to incorporate habitat use models that are shared by popular inferential approaches such as Maximum Entropy^57,58^ and Resource Selection Functions^59,60^. Habitat selection originates from the idea of disproportionate use, compared to the availability of a habitat. Therefore, a habitat selection function is defined in terms of the ratio of usage per unit of habitat available

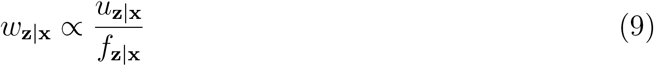

This expression is most often encountered in its unconditional form *w*_**z**_ ∝ *u*_**z**_/*f*_**z**_. Biologically, an unconditional formulation means that there are no mobility constraints and the animal has access to any habitat in proportion to its availability across the entire landscape^60^. Eq. 9 implies a definition for conditional habitat usage^61^.

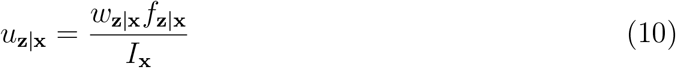

Where the denominator is a normalizing integral *I*_**x**_ = ∫_*E*_ *w*_**z**|**x**_*f*_**z**|**x**_*d***z**. Animal mobility and behaviour can complicate our formulations of both of the two main determinants of usage in eq. (7), so the notions of habitat availability and habitat preference must be re-evaluated in the light of animal mobility.

### 2.3 Mobility and habitat availability

The simplicity of the definition of habitat availability *f*_**x**_ as the amount (e.g. total area) of a habitat that is accessible to a population is deceptive, because it does not easily yield to quantification for a given species in a given landscape. This makes the definition of habitat availability one of the most challenging and influential attributes of species-habitat association models. As shown by a number of earlier studies^56,62,63^, the quantitative representation of what is available to animals can alter the parameter estimates and predictions of such models, often converting underlying preference to apparent avoidance of particular habitats, and vice-versa. Two aspects of movement in particular require attention. The first refers to accessibility of any point in *E*-space from another, and the second relates to complementary use of habitats by means of commuting.

Dealing with accessibility between habitats **z** and **x**, requires us to port geographical measures such as mobility and spatial autocorrelation into *E*-space. However, to manipulate habitat availability mathematically, we first need to represent it parametrically. Habitat availability across an arbitrarily large geographical domain, can be approximated in *n* environmental dimensions by a Gaussian mixture of *L* components^35^

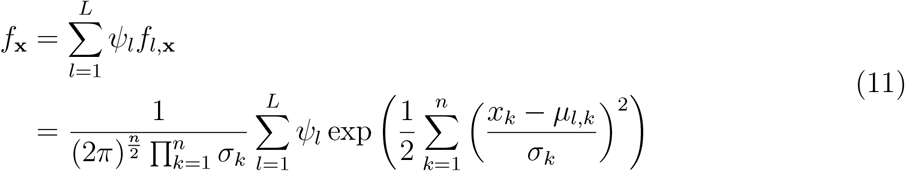

where *f_l,_*_**x**_ is the *l^th^* mixture component (a unimodal probability density function in *n* dimensions), *ψ_l_* is the weight associated with the *l^th^* component (such that 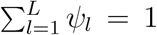), *μ_l,k_* is the mean (i.e. the location in *E*-space) of the *l^th^* mixture component along the *k^th^* environmental dimension and *σ_k_* is the characteristic standard deviation along the *k^th^* environmental dimension. This form of unconditional availability gives us the probability density of any given habitat across the whole of space. This Gaussian mixture approximation is not the only way to capture *n*-dimensional hypervolumes, but other approaches are similar in spirit^30^. We can extend these ideas to define conditional availability *f* _**z**|**x**_ which describes the frequency with which different habitats **z** would be accessible close to a reference habitat **x** and depends on organism mobility and environmental autocorrelation. Recently, an expression was derived^33^ for conditional habitat availability for orthogonal environmental variables (i.e. either raw environmental variables presenting no cross-correlation, or rotated covariates via a method such as principal components analysis), as perceived from the vantage point of an organism found at habitat **x**.

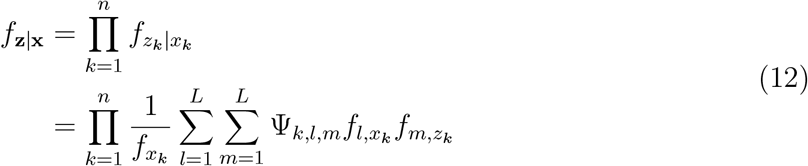

where, as in eq. (11), 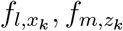 are respectively the *l^th^* and *m^th^* Gaussian components for the *k^th^* environmental dimension at the values *x, z*. The Gaussian mixture 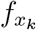 at the value *x* of the *k^th^* variable is calculated using the new weights Ψ_*k,l,m*_ that are derived as a function of the mobility of the study organism combined with empirical curves of the spatial autocorrelation of the environmental covariates (see Appendices in 33).

A feature not yet fully explored by these frameworks is complementarity in habitat use, which leads to the “zero-niche paradox” mentioned in the Introduction. If an organism can commute between two habitats, it can use their properties in a complementary way. For example, a habitat that provides water and one that provides food may be insufficient for survival, on their own, but entirely adequate to support an organism when used in combination. By commuting, an animal effectively creates a third, sufficient habitat (containing both water and food). This can be thought of as the capacity of mobile organisms to alter habitat availability, depending on spatial context, i.e. the total set of habitats that physically exist and are within reach of an organism^13^. The above approach, constructing Gaussian approximations of observed availabilities, has the potential to capture complementarity by smoothing these observed frequencies into approximate probability densities. Through this smoothing operation, habitats that do not physically exist, but are proximate to extant habitats in *E*-space, receive a non-zero density. This effectively makes them available to the animals. Although the details of how best to achieve this smoothing to faithfully capture complementarity between habitats are an open research question, it is important to stress the expandability of the approach presented here.

### 2.4 Mobility and habitat preference

Following the requisites of HSF frameworks such as RSFs and MaxEnt, habitat preference is broadly expressed as an exponential transformation of a predictor function *g*(**x**).

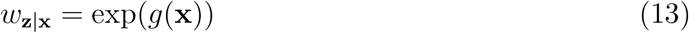

Echoing the formulation for unconditional fitness in eq. (8) the predictor function *g* (**x**) can be formulated as a 2nd order polynomial in the dimensions of the vector **x** and for some coefficients *γ*(note that the coefficients *γ_r,k_* of habitat preference are not the same as the fitness coefficients *β_r,k_* in eq. (8)).

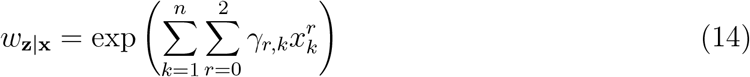

There is an extensive literature describing how the use of a particular habitat can be affected non-linearly by the availability of surrounding habitats, a phenomenon called a functional response in habitat selection^64,65^. To resolve this, it was suggested^59^ that functional responses could be flexibly captured by expressing the *γ* coefficients of eq. (14) as functions of the entire habitat availability field *γ*(**f**). In its simplest form, this may be written as a linear combination of the availabilities of all habitats across *E*-space:

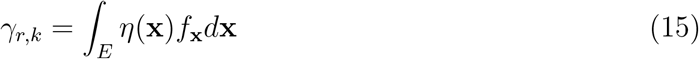

For some function *η* that describes how the coefficient *γ* responds to the availability of any given habitat **x**. This varying-coefficients approach was named a *Generalized Functional Response* (GFR)^32^ and a particular version of GFRs was formulated, by expressing the *η*’s as polynomial functions of environmental variables. This is the simplest formulation of GFRs and leads to an expression for the coefficients of habitat preference, in terms of the moments of the marginal distributions of habitat availability along each environmental dimension

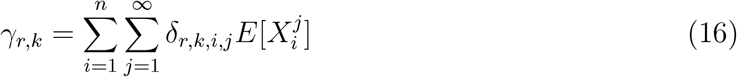

In practical applications, due to limitations in data availability, only the lower moments are used (i.e. the average value of each environmental variable in the neighborhood of the point of interest).

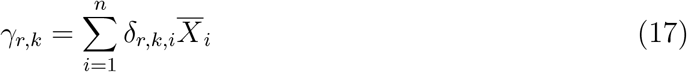

The exploration of efficient (i.e. economical with degrees of freedom) and effective (i.e. accurate and precise) GFR models is still at its early stages and improvements of implementing the general idea of eq. (15) will be possible. However, currently the state-of-the-art with GFRs is eq. (16), and this has repeately been able to improve the predictive abilities of HSF models^32,35,36,56,63,66^.

### 2.5 Parametric definition of the fundamental niche

The condition in eq. (6) can now be expanded with the aid of eqs (7) and (10)

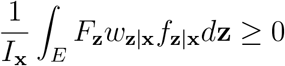

Given that the normalizing constant *I*_**x**_ is consistently non-negative, we can simplify the above expression into an equation for the boundary of the fundamental niche

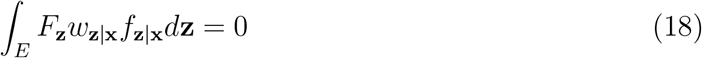

The three quantities participating in the integral are estimable from field data. The habitat-specific fitness *F*_**z**_ has been formulated mathematically^35^ and fitted to space use and population growth data from the field^36^.This framework is a specific example of the much broader class of hybrid models proposed conceptually by^26^. The habitat preference model *w*_**z**|**x**_, once recast as a GFR model^32^ can be estimated from usage data^32,36^. Finally, the conditional availability can be derived from a universal approximator, such as the Gaussian mixture model^33,35^. Eq. (18) relies on different categories of parameters. The function *F*_**z**_ contains the vector of *fitness parameters β*, the function *w*_**z**|**x**_ contains the vector of *habitat use parameters γ* and the function *f*_**z**|**x**_ contains the *habitat availability* parameters, which in the case of Gaussian mixtures would principally be (*μ, σ*), the locations and variances of the Gaussian mixture components. There may be other, more general formulations for the participating functions in eq. (18) and their respective parameters. These may broaden the applicability of this very general framework to a wealth of life-histories for different animals. For some biological scenarios, it is already possible to specify these functions and proceed with particular mathematical formulations of the niche.

### 2.6 The fundamental niche of house sparrows

By focusing on colonial or territorial species, it is possible to use a single definition of habitat availability for each sampling instance (i.e. the domain of each colony, or the territory of each individual or group). This definition of the sampling instance, means that a simple and distinct Gaussian mixture in eq. (11) can be used to describe available habitats for each instance, instead of the conditional approxiation of eq. (12).

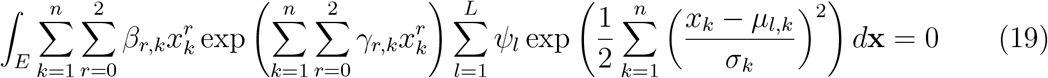

This expression has a closed form (see Appendix A in^35^).

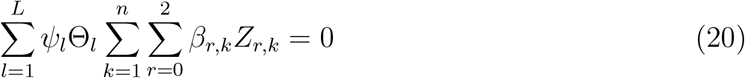

where

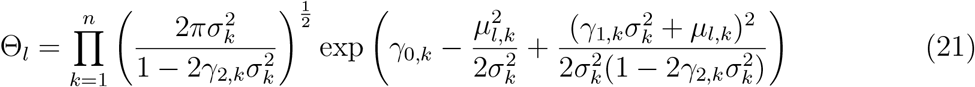

and

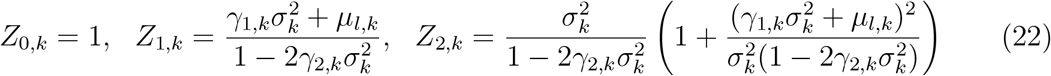

Note that this expression is independent of **x**. It depends purely on the fitness, habitat use and habitat availability parameters. Given that the fitness parameters are fixed (independent of habitat) and that habitat use parameters depend on habitat availability (via a GFR), then the fundamental niche in this example depends on the habitat composition of the home range of the individuals.

The habitat selection model developed for the sparrow data in^36^ contained no quadratic terms in any of the habitat variables, which implies that *γ*_2,*k*_ = 0 for all *k*. This greatly simplifies eqs (21) and (22) to

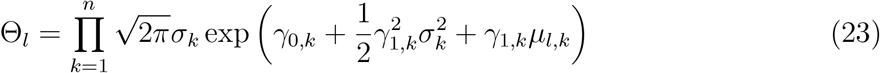

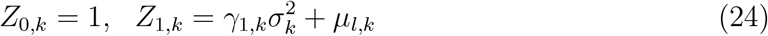

The expression for *Z*_2,*k*_ is ommitted because, by definition *β* _2,*k*_ = 0. The parameters of this expression are data-driven^36^, but the model can be asked to describe the niche in simplified versions of environmental space, allowing mathematical tractability and revealing some highly informative features of the niche. To begin, let us assume that the habitat availability within a home range of a sparrow colony can be described by a single (i.e. *L* = 1 and _1_ = *ψ*_1_) Gaussian component of *n* orthogonal variables (e.g. 67), so that *f*_**x**_ = *MN* (*μ, σ***I**_*n*_), where *μ* = (*μ*_1_*, …, μ_n_*) are the average values of each environmental variable, *σ* is a shared standard deviation and **I**_*n*_ is the *n* × *n* identity matrix. The shared *σ* will be used here as a convenient shorthand for environmental heterogeneity (as *σ* → 0, the entire home range comprizes identical cells, each with composition *μ*). These simplifications give a very tractable version of the niche for sparrows

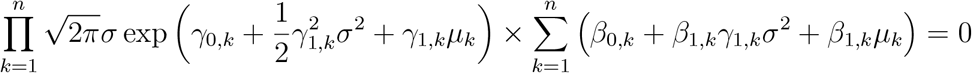

Given that the first component of this product can never be zero, the expression further reduces to

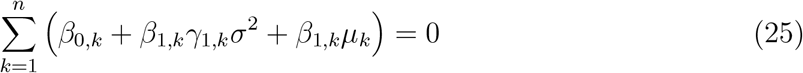

Under the GFR framework, the habitat selection coefficients can be written as functions of moments from the marginal distributions of the environmental variables. In^36^ these were sufficiently modelled as linear functions of the averages of all covariates, so that 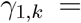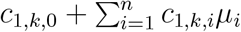. This affords an expression for the niche in terms of purely *β, σ, μ*

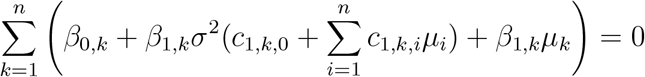

Defining 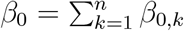

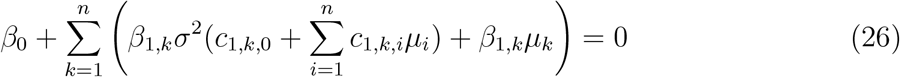

This expression was used to explore the fundamental niche of sparrows, within the much simplified, unimodal version of *E*-space. Given any position for the average habitat composition **x** = (*μ_R_, μ_G_, μ_B_*), for roof, grass and bush respectively, only points that satisfied *μ_R_*+*μ_G_*+*μ_B_* ≤ 100% were considered. Then, the variability of habitats around that point was examined by increasing the value of *σ* to its maximum. The maximum value of *σ* depends on **x**. To preserve the 100% summation constraint for proportions in habitat units within the sparrow home range, I required that 95% of the density of the availability sphere in *E*-space would fall within the feasible region. Note that the three land cover variables were not exhaustive, because the habitat was characterized by other covariates for land cover. So, their sum could be any number between The maximum radius of a sphere from the feasibility boundaries of the niche space is *r_max_* = min(*μ_R_, μ_G_, μ_B_, d*), where 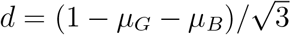 is the distance of the centre-point from the inclined plane of 100% land cover. We therefore require that 95% of the probability density contour of a spherical Gaussian would be at that distance from the centre. For a multivariate normal *N* (**0***, σI*_3_), the square of the Mahalanobis distance is chi-square distributed, with critical value at 95% of 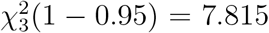. So, the critical distance is 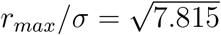. This implies that the maximum value *σ_max_* of *σ* is 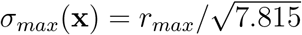. Within the interval *σ* ∈ [0*, σ_max_*(**x**)], I defined viability as the occurrence of a positive value on the left-hand-side of eq. (25), for the homogeneous scenario, and eq. (26) for the heterogeneous scenario. I also recorded these values as the measure of fitness at each of reference habitat **x**. For the homogeneous scenario, only one value of viability and fitness corresponded to eaxh reference habitat. For the heterogeneous scenario, I recorded the maximum achievable fitness and the corresponding value of habitat heterogeneity *σ* that produced it.

I generated the following statistics to examine the impact of heterogeneity and habitat selection on the niche. First, the percentage of feasible environmental space that contained viable reference habitats for the homogneous case. Second, the percentage of homogeneous habitats whose fitness was improved with the addition of heterogeneity and selection. Third, the percentage of inviable habitats that were made viable with the addition of heterogeneity and selection. Fourth, the percentage of increase in fitness resulting from the addition of heterogeneity and selection.

## References

1. Mcinerny, G. J. & Etienne, R. S. Ditch the niche - is the niche a useful concept in ecology or species distribution modelling? Journal of Biogeography 39, 2096–2102 (2012).

2. Hutchinson, G. Concluding remarks. in Cold spring harbor symposia on quantitative biology 75–96 (1957). doi:10.1201/9781315366746.

3. Carscadden, Kelly, A. et al. Niche Breadth: Causes and consequences for Ecology, evolution, and conservation. 95, 179–214 (2020).

4. Pocheville, A. The ecological niche: History and recent controversies. in Handbook of evolutionary thinking in the sciences 547–586 (2015). doi:10.1007/978-94-017-9014-7_26.

5. Chase, J. M. & Leibold, M. A. Ecological Niches: Linking Classical and Contemporary Approaches. 212 (Springer Science+ Business Media BV, Formerly Kluwer Academic Publishers BV, 2003).

6. Peterson, A. T. et al. Ecological niches and geographic distributions. vol. 56 314 (Princeton University Press, 2011).

7. Colwell, R. K. & Fuentes, E. R. Experimental sudies of the niche. Annual Review of Ecology, Evolution, and Systematics 6, 281–310 (1975).

8. Pulliam, H. R. On the relationship between niche and distribution. Ecology Letters 3, 349–361 (2000).

9. Chesson, P. A need for niches? Trends In Ecology & Evolution 6, 26–29 (1991).

10. Araújo, M. B. & Guisan, A. Five (or so) challenges for species distribution modelling. Journal of Biogeography 33, 1677–1688 (2006).

11. Mcinerny, G. J. & Etienne, R. S. Stitch the niche - a practical philosophy and visual schematic for the niche concept. Journal of Biogeography 39, 2103–2111 (2012).

12. Mcinerny, G. J. & Etienne, R. S. Pitch the niche - taking responsibility for the concepts we use in ecology and species distribution modelling. Journal of Biogeography 39, 2112–2118 (2012).

13. Angilletta, M. J., Sears, M. W., Levy, O., Youngblood, J. P. & VandenBrooks, J. M. Fundamental Flaws with the Fundamental Niche. Integrative and Comparative Biology 59, 1038–1048 (2019).

14. Warren, D. L. In defense of ‘niche modeling’. Trends in Ecology and Evolution 27, 497–500 (2012).

15. Kearney, M. R. & Porter, W. P. Mapping the fundamental niche: Physiology, Climate and the distribution of a nocturnal lizard. Ecology 85, 3119–3131 (2004).

16. Soberón, J. Commentary on Ditch, Stitch and Pitch: The niche is here to stay. Journal of Biogeography 41, 414–417 (2014).

17. Yates, K. L. et al. Outstanding Challenges in the Transferability of Ecological Models. Trends in Ecology and Evolution 33, 790–802 (2018).

18. Soberón, J. & Peterson, A. T. What is the shape of the fundamental Grinnellian niche? Theoretical Ecology 13, 105–115 (2020).

19. Godsoe, W., Jankowski, J., Holt, R. D. & Gravel, D. Integrating Biogeography with Contemporary Niche Theory. Trends in Ecology and Evolution 32, 488–499 (2017).

20. Holt, R. D. Bringing the Hutchinsonian niche into the 21st century: Ecological and evolutionary perspectives. Proceedings of the National Academy of Sciences of the United States of America 106, 19659–19665 (2009).

21. Godsoe, W. I can’t define the niche but i know it when i see it: A formal link between statistical theory and the ecological niche. Oikos 119, 53–60 (2010).

22. McInerny, G. J. & Etienne, R. S. ‘Niche’ or ‘distribution’ modelling? A response to Warren. Trends in Ecology and Evolution 28, 191–192 (2013).

23. Whittaker, R. H., Levin, S. A. & Root, R. B. Niche, Habitat and Ecotope. The American Naturalist 107, 321–338 (1973).

24. Soberón, J. & Nakamura, M. Niches and distributional areas: Concepts, methods, and assumptions. Proceedings of the National Academy of Sciences of the United States of America 106, 19644–19650 (2009).

25. Elith, J. & Leathwick, J. R. Species distribution models: ecological explanation and prediction across space and time. Annual Review of Ecology, Evolution, and Systematics 40, 677 (2009).

26. Schurr, F. M. et al. How to understand species’ niches and range dynamics: A demographic research agenda for biogeography. Journal of Biogeography 39, 2146–2162 (2012).

27. Blonder, B. Hypervolume concepts in niche- and trait-based ecology. Ecography 41, 1441–1455 (2018).

28. Holt, R. D. On the Relation between Niche Overlap and Competition : The Effect of Incommensurable Niche Dimensions. Nordic Society Oikos 48, 110–114 (1987).

29. Malanson, G. P. Simulated responses to hypothetical fundamental niches. Journal of Vegetation Science 8, 307–316 (1997).

30. Blonder, B., Lamanna, C., Violle, C. & Enquist, B. J. The n-dimensional hypervolume. Global Ecology and Biogeography 23, 595–609 (2014).

31. Blonder, B. Do hypervolumes have holes? American Naturalist 187, E93–E105 (2016).

32. Matthiopoulos, J., Hebblewhite, M., Aarts, G. & Fieberg, J. Generalized functional responses for species distributions. Ecology 92, 583–589 (2011).

33. Matthiopoulos, J., Fieberg, J., Aarts, G., Barraquand, F. & Kendall, B. E. Within reach? Habitat availability as a function of individual mobility and spatial structuring. American Naturalist 195, 1009–1026 (2020).

34. McGill, B. J. Matters of scale. Science 328, 575–576 (2010).

35. Matthiopoulos, J. et al. Establishing the link between habitat selection and animal population dynamics. Ecological Monographs 85, 413–436 (2015).

36. Matthiopoulos, J., Field, C. & MacLeod, R. Predicting population change from models based on habitat availability and utilization. Proceedings of the Royal Society B: Biological Sciences 286, (2019).

37. Monahan, W. B. & Tingley, M. W. Niche tracking and rapid establishment of distributional equilibrium in the house sparrow show potential responsiveness of species to climate change. PLoS ONE 7, (2012).

38. Peterson, A. T., Papeş, M. & Soberón, J. Mechanistic and correlative models of ecological niches. European Journal of Ecology 1, 28–38 (2015).

39. Guisan, A. & Zimmermann, N. E. Predictive habitat distribution models in ecology. Ecological Modelling 135, 147–186 (2000).

40. Kissling, W. D. et al. Towards novel approaches to modelling biotic interactions in multispecies assemblages at large spatial extents. Journal of Biogeography 39, 2163–2178 (2012).

41. Ovaskainen, O. & Abrego, N. Joint Species Distribution Modelling: With Applications in R (Ecology, Biodiversity and Conservation). 388 (Cambridge University Press, 2020).

42. Leibold, M. A. Ecology: Return of the niche. Nature 454, 39–41 (2008).

43. Hall, L. S., Krausman, P. R. & Morrison, M. L. The Habitat Concept and a Plea for Standard Terminology. Wildlife Society Bulletin 25, 173–182 (1997).

44. Aarts, G., MacKenzie, M., McConnell, B., Fedak, M. & Matthiopoulos, J. Estimating space-use and habitat preference from wildlife telemetry data. Ecography 31, 140–160 (2008).

45. Turchin, P. Complex population dynamics: A theoretical/empirical synthesis. 450 (Princeton University Press, 2003).

46. Stenseth, N. C. Grasses, grazers, mutualism and coevolution: A comment about hand-waving in ecology. Oikos 41, 152–153 (1983).

47. Nur, N. Fitness, population growth rate and natural selection. Oikos 42, 413–414 (1984).

48. Murray, B. G. Population Growth Rate as a Measure of Individual Fitness. Oikos 44, 509 (1985).

49. Nur, N. Population Growth Rate and the Measurement of Fitness: A Critical Reflection. Oikos 48, 338 (1987).

50. Ollason, J. G. What Is This Stuff Called Fitness ? Biology and Philosophy 6, 81–92 (1991).

51. Fisher, R. The genetical theory of natural selection. (The Clarendon Press, 1930).

52. Lande, R. A quantitative genetic theorey of life history evolution. Ecology 63, 607–615 (1982).

53. Roff, D. A. Defining fitness in evolutionary models. Journal of Genetics 87, 339–348 (2008).

54. Matthiopoulos, J. The use of space by animals as a function of accessibility and preference. Ecological Modelling 159, 239–268 (2003).

55. Austin, M. Species distribution models and ecological theory: A critical assessment and some possible new approaches. Ecological Modelling 200, 1–19 (2007).

56. Paton, R. & Matthiopoulos, J. Defining the scale of habitat availability for models of habitat selection. Ecology 97, 1113–1122 (2018).

57. Elith, J. et al. A statistical explanation of {MaxEnt} for ecologists. Diversity and Distributions 17, 43–57 (2011).

58. Merow, C., Smith, M. J. & Silander, J. A. A practical guide to MaxEnt for modeling species’ distributions: what it does, and why inputs and settings matter. Ecography 36, 1058–1069 (2013).

59. Boyce, M. S., McDonald, L. L. & Manly, B. F. J. Relating populations to habitats-Reply. Trends In Ecology & Evolution 14, 490 (1999).

60. Manly, B. F., Thomas, D., McDonald, T. L. & Erickson, W. P. Resource Selection by Animals. 222 (Springer Netherlands, 2002). doi:10.1007/0-306-48151-0.

61. Lele, S. R. & Keim, J. L. Weighted distributions and estimation of resource selection probability functions. Ecology 87, 3021–3028 (2006).

62. Beyer, H. L. H. L. et al. The interpretation of habitat preference metrics under use– availability designs. Philosophical Transactions of the Royal Society of London B: Biological Sciences 365, 2245–2254 (2010).

63. Aarts, G., Fieberg, J., Brasseur, S. & Matthiopoulos, J. Quantifying the effect of habitat availability on species distributions. Journal of Animal Ecology 82, 1135–1145 (2013).

64. Mysterud, Atle; Ims, R. A., Mysterud, A. & Ims, R. A. Functional Responses in Habitat Use : Availability Influences Relative Use in Trade-Off Situations. Ecology 79, 1435–1441 (1998).

65. Holbrook, J. D. et al. Functional responses in habitat selection: clarifying hypotheses and interpretations. Ecological Applications e01852 (2019) doi:10.1002/eap.1852.

66. Muhly, T. B. et al. Functional response of wolves to human development across boreal North America. Ecology and Evolution 9, 10801–10815 (2019).

67. Jiménez, L., Soberón, J., Christen, J. A. & Soto, D. On the problem of modeling a fundamental niche from occurrence data. Ecological Modelling 397, 74–83 (2019).

